# Prediction of misfolded proteins spreading in Alzheimer’s disease using machine learning

**DOI:** 10.1101/2022.10.04.510701

**Authors:** Luca Gherardini, Aleksandra Pestka, Lorenzo Pini, Alessandro Crimi

## Abstract

The pervasive impact of Alzheimer’s disease on aging society represents one of the main challenges at this time. Current investigations highlight two specific misfolded proteins in its development: Amyloid-*β* and *τ*. Previous studies focused on spreading for misfolded proteins exploited simulations, which required several parameters to be empirically estimated. Here, we provide an alternative view based on a machine learning approach. The proposed method applies an autoregressive model, constrained by structural connectivity, to predict concentrations of Amyloid-*β* two years after the provided baseline. In experiments, the autoregressive model generally outperformed the state-of-art models yielding the lowest average prediction error (mean-squared-error 0.0062). Moreover, we assess its effectiveness and suitability for real case scenarios, for which we provide a web service for physicians and researchers. Despite predicting amyloid pathology alone is not sufficient to clinical outcome, its prediction can be helpful to further plan therapies and other cures.

## Introduction

The study of the mechanisms underlying neurodegenerative diseases is extremely important to address the epochal challenges we are facing, where world populations are aging rapidly, and dementia incidence is increasing sharply. Alzheimer’s disease (AD) is the most common form of neurodegeneration. To date, no cure is available for this disease. Recent significant advancement in the pharmacological field has been done, although findings are still far from being conclusive (*1–3*). A better understanding of the pathophysiological mechanisms underlying AD, jointly with non-invasive imaging biomarkers, might pave the way to novel pharmacological and non-pharmacological treatments. Recent advancements in magnetic resonance imaging (MRI) have provided better estimations of brain properties through in-vivo analysis (*4,5*), allowing a deeper understanding of the evolution of AD neuropathological changes (NC). Diffusion-weighted imaging (DWI) shows the information on the anatomical connectivity inside the brain, polarizing water molecules to detect their diffusion in tissues to identify white matter tracts through tractography. The structural connectome (*6*) represents the brain network using regions provided by an anatomical atlas (*4*) as nodes and white fibers as weighted edges, exploiting graph theory to obtain a more advanced understanding of the mechanics based on structural connectivity.

In AD, the diffusion pattern revolves mainly around two misfolded proteins (MP), namely *Amyloid* − *β* (*Aβ*) and tau (*τ*), across well-known pathways (*7*). According to the “molecular nexopathy” model, large-scale networks might provide the biological substrates for the spreading of MP, producing macroscopic signatures of network dysfunction (*8*).

This assumption allows the prediction of future trajectories and the speed of progression, alternatively recent research drawing specific diffusion patterns related to genetic and cerebrospinal fluid (CSF) biomarkers (*9*). Recent views (*10*) consider AD as a two-stage disease, where *Aβ* has the heavier effect in the first disease stage, preceding and driving the onset of clinical impairment for years, before being shadowed by *τ* at the display of symptoms (*11*).

Notably, these stages have differential effects on brain network connectivity (*10*). Nevertheless, their spatio-temporal trajectories are discordant, *Aβ* might even slow down at later stages, conversely to *τ* which steadily keeps increasing for the remaining course of AD (*11*). Recently, Collij et al. (*12*) question a single-universal trajectory of *Aβ* progression reporting the existence of different subtypes of topographic *Aβ* accumulation. This is in line with the recent probabilistic framework of AD. While the “classical amyloid hypothesis” states that *Aβ* deposition is a causative agent of AD, being sufficient and necessary, Frisoni and colleagues (*13*) proposed a probabilistic model in which *Aβ* is still a key player in AD-NC, but the penetrance of the amyloid cascade is directly proportional to the penetrance of genetic risk factors. Based on these premises, it is crucial to map the spreading of those volatile and variable MPs to follow and tackle the initial progression, and to identify those subjects who will remain stable or would show specific trajectories. Through radiopharmaceutical tracers, it is possible to obtain positron emission tomography (PET) images to measure the spatio-temporal cerebral concentrations of proteins. AD, Parkinson’s disease (PD), and amyotrophic lateral sclerosis have also been described in other studies by mathematical models. Those range from epidemic-based spreading on the connectome (*14*) to an analytical model of protein movement based on the heat equation under a connectivity-driven mechanism (*15*). A more recent computational approach predicts how those misfolded proteins interact with receptors, simulated regional neural activity, and electroencephalograms (*16*). Other studies investigated connectivity based on co-variability of PET tracers (*17–19*). A recent review summarized these findings and other connectomics related studies (*20*).

Given the current trend motivated by exceptional results with machine learning methods in many fields, we adopted a different approach, inspired by a model used to predict bloodoxygen-level-dependent (BOLD) signals constrained by tractography (*21*). This is motivated by the hope of finding more accurate results. Moreover, even assuming comparable efforts in parameters fine tuning. Simulation based approaches are dependent on the design of a model, while a purely data-driven approach like the one proposed will not require such a complicated design, as mentioned in other machine learning versus simulation-based comparison (*22*). However, it is dependent on the available data.

More specifically, we predicted misfolded protein depositions using an auto-regressive multivariate model (MAR) given baseline concentrations in each region of interest (ROI), and the structural connectome as a physical constraint for the spreading, limiting the solution space. Even though *τ* has a stronger correlation with advanced cognitive decline, in this paper we focus on *Aβ* as we are looking not only at AD patients at the beginning of the disease but also MCI. Nevertheless, the approach should work also with different misfolded proteins. The approach is summarized in figure 1. We also developed a graph convolutional network (GCN) to apply a different learning approach to the natural graph representation of such phenomena, as an alternative still from the machine learning perspective.

**Figure 1:**
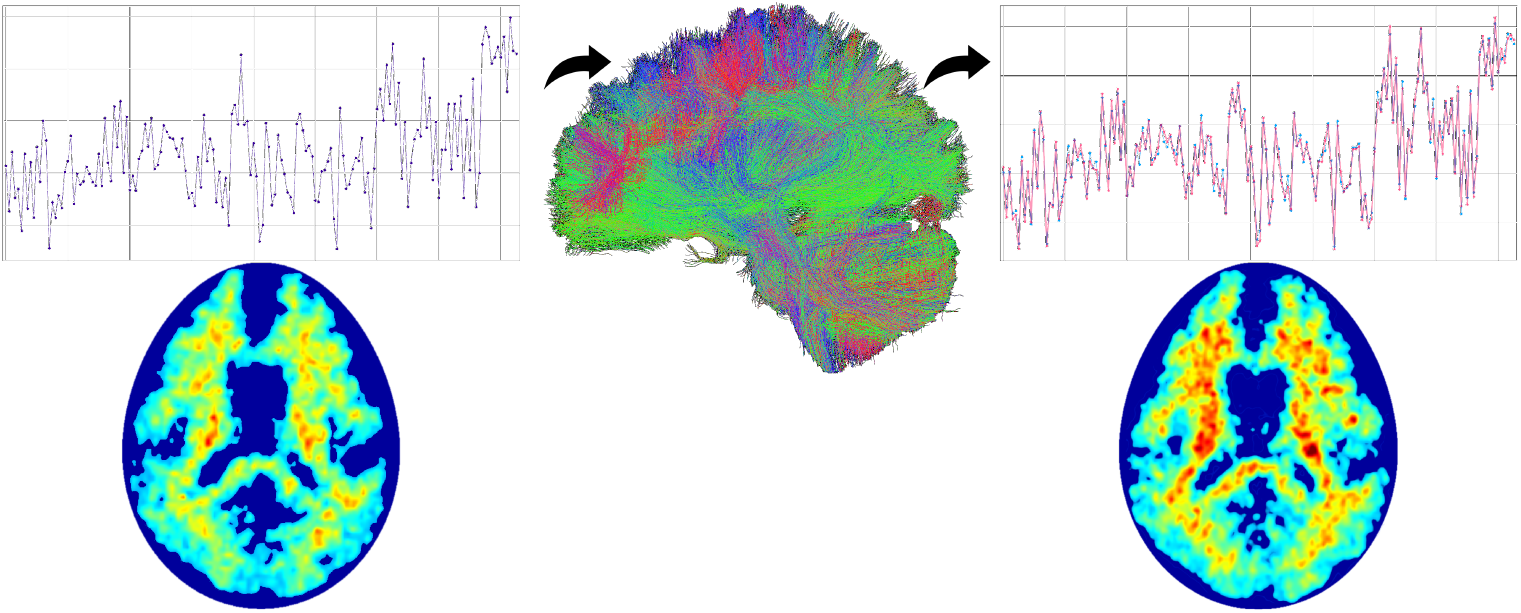
Idealized representation of the approach. Given a baseline misfolded protein regional deposition and a structural connectome, a future protein deposition is predicted.

We tested the approach in a cross-validation manner on the publicly available ADNI dataset (https://ida.loni.usc.edu) for a total of 212 subjects. Details on the selection of subjects, preprocessing, and simulation settings are available in the Methods section.

The performance evaluation gauged the reliability of predicted deposits using follow-up PET as ground truth (in our experiments 24 months, though longer periods are possible). The validity of the proposed approach challenged two state-of-the-art models considered in this scenario, respectively the *Epidemic Spreading Model* (ESM) (*14*) and the *Network Diffusion Model* (NDM) (*15*), and statistics scrupulously gathered allowed to debate the winner and the assets of each model. Lastly, we validated the accessibility of the proposed tool offering the prediction as an online service for collaborative brain research. The proposed predictive model is available at https://brainspread.sano.science/ where users can upload a structural connectome and a PET file from the same subject to predict misfolded protein depositions. This new modeling framework could have significant clinical implications, helping to predict the evolution of A spreading in patients with early clinical manifestations, useful for enrichment of AD clinical trial at the predementia stage, thus with individuals with clinical impairment and likely to progress rapidly.

We also exploited the MAR to predict clinical dementia ratio (CDR) scores, available on the ADNI database, from PET images and CDR scores at baseline.

## Results

We performed the simulations on both distinct clinical categories (i.e., AD, mild cognitive impairment (MCI), and controls (CN)) and the whole dataset/cohort, following the modalities described in the *Methods* section. MAR and GCN used a leave-one-out cross-validation approach. We reported, with each configuration, its computational time using ten cores on an Intel i9-11900KF CPU. Each simulation produced a report with its statistics, especially the average mean squared error (MSE) and Pearson’s correlation coefficient (PCC), reported in tables and individual values. In figure 2 is it possible to observe the predictions for an exemplative subject.

**Figure 2:**
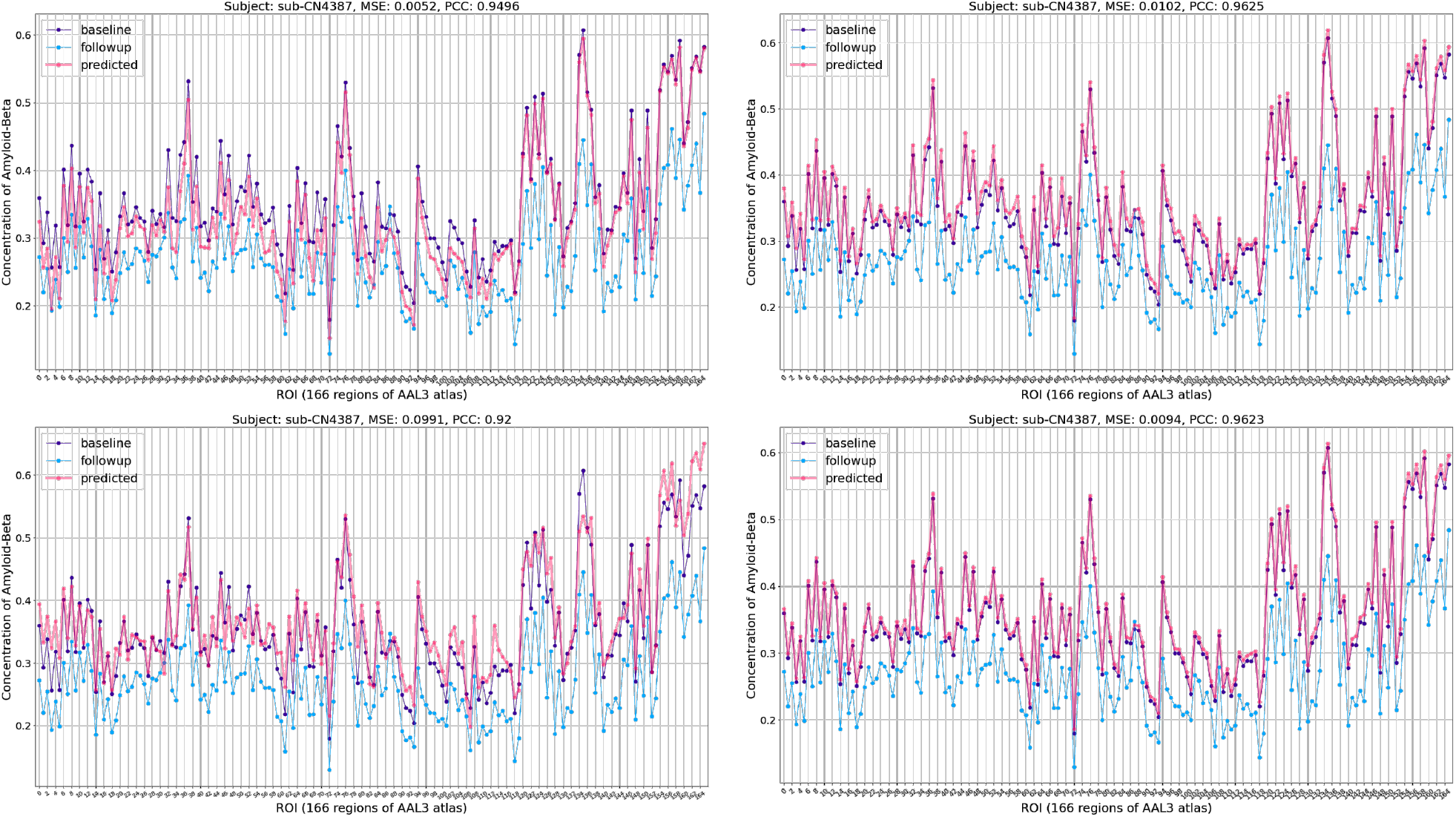
Plots reporting predictions for the CN subject #4387 by MAR (upper left), ESM (upper right), GCN (bottom left) and NDM (bottom right). This is an example of decaying Amyloid-*β* that is handled correctly by MAR but not by ESM and NDM simulations, which always go for incremental predictions.

CDR prediction of MAR tended to give values around 0.5, showing a normal distribution, as in figure 3. The Mann-Whitney U test confidently confirmed the tendency to assign lower CDRs, with a p-value of 4.66e-04.

**Figure 3:**
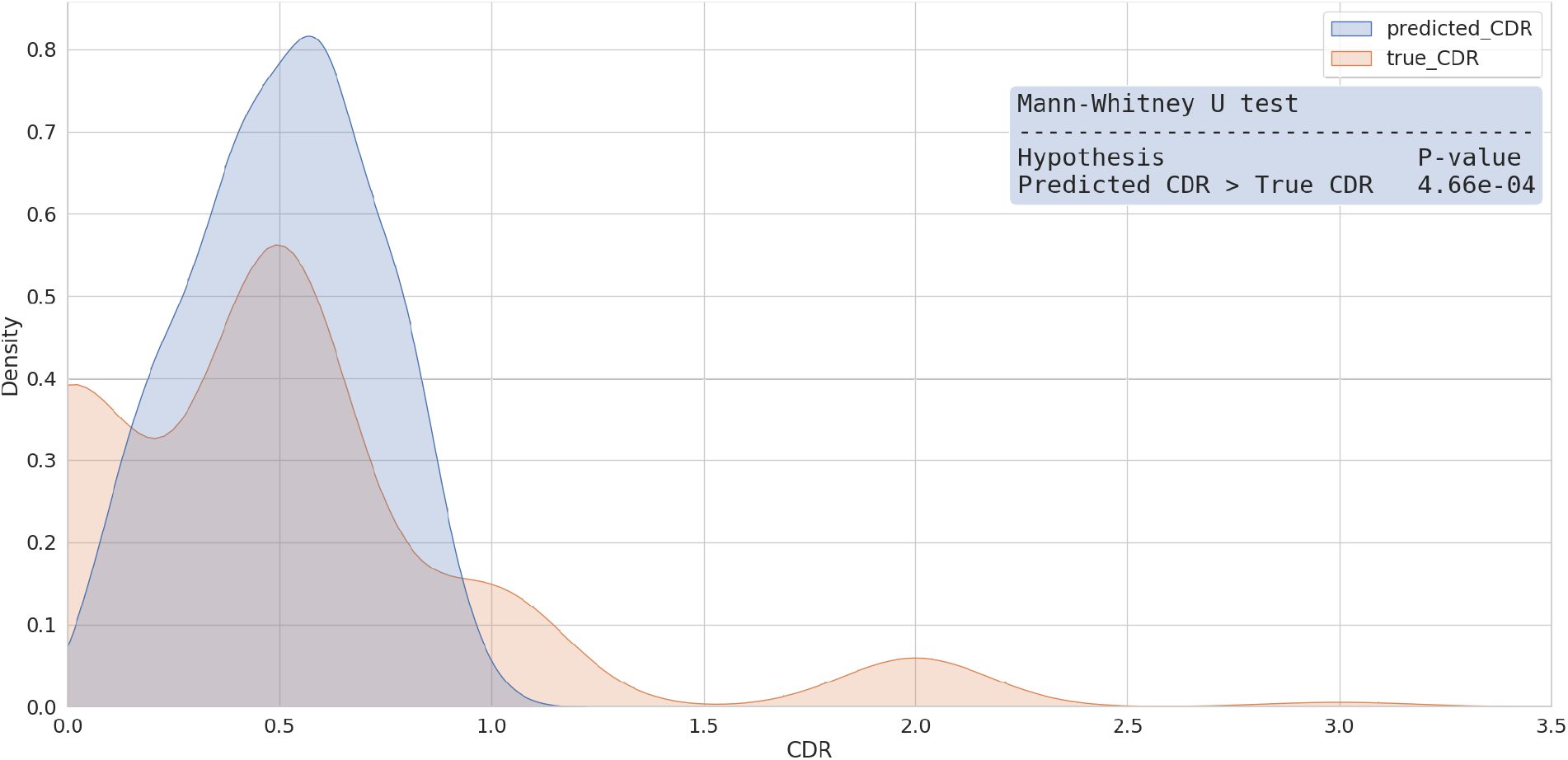
Plot displaying the density of predicted and true CDR scores from the alternative MAR model (left). The Mann-Whitney U test (right) rejected the alternative hypothesis that the predicted CDRs were statistically higher than the true ones with a p-value of 4.66e-04.

### Regional error

We additionally analyzed the regional error made by these models in their best configuration and on the whole dataset/cohort, displaying their location through Brain Net Viewer (*23*) in figure 4. Looking at the distribution of regional errors in the lower plot of figure S1, it is observable a considerable shift towards the right for the GCN distribution, and the ESM has a right-skew longer than MAR, suggesting the tendency to make weightier errors. The Mann-Whitney U test, displayed in figure S1, confirms these two hypothesis, but doesn’t remain under the 95% confidence interval with the NDM.

**Figure 4:**
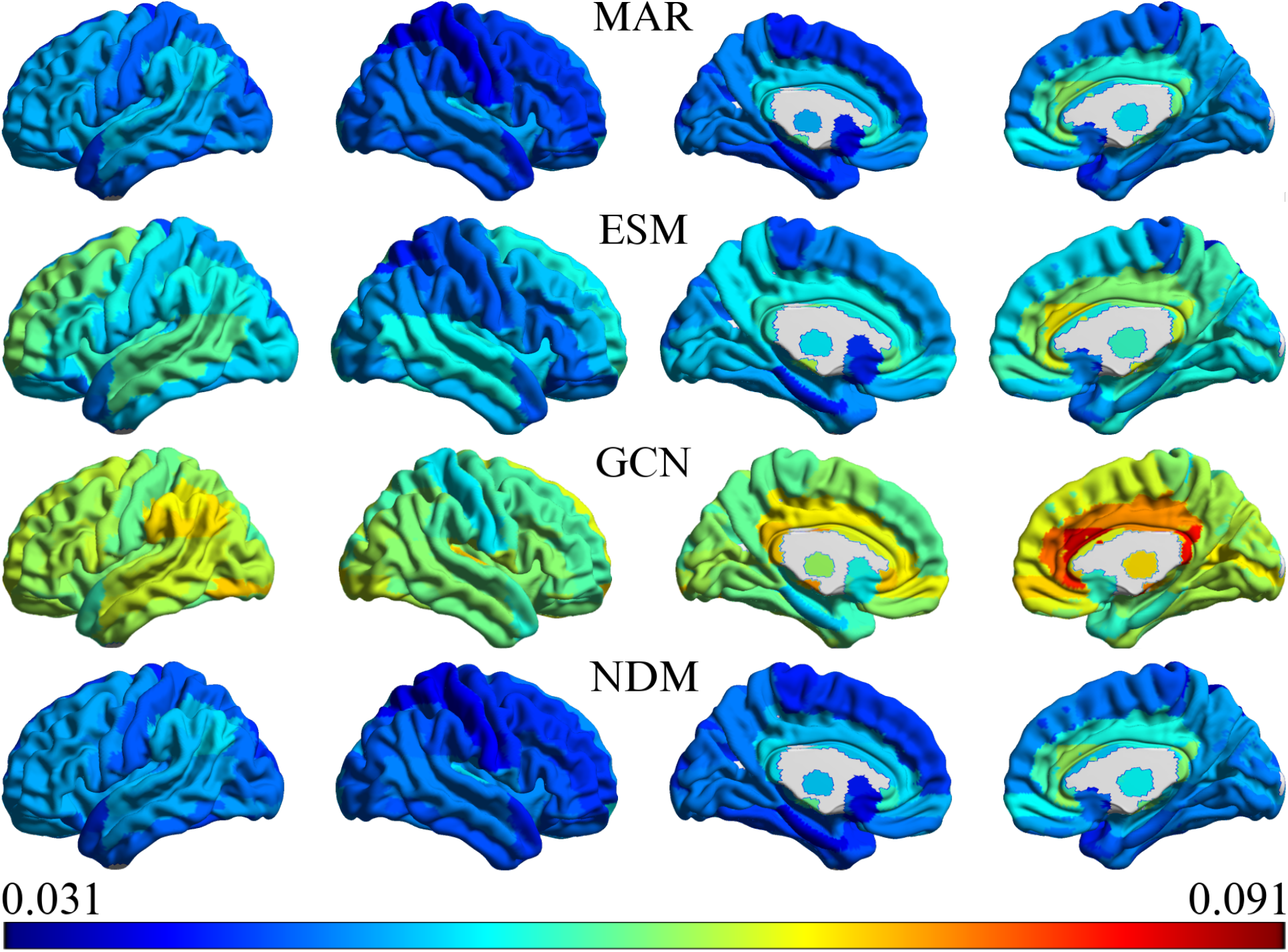
Mapping, realized with Brain Net Viewer (*23*), of the regional errors for the best configurations of MAR (first), ESM (second), GCN (third), and NDM (fourth row). The scale comprises the minimum and maximum values across all four. Interestingly, the regional error appears very low for MAR and NDM in the *somatory sensor motor cortex* (third and fourth column), while ESM and GCN showed higher error in the medial and lateral temporal cortex, key hubs of Amyloid-*β* deposition in Alzheimer’s disease.

Subjects with no increase or partial decrease of *Aβ*

In the dataset we considered, some subjects spanning all clinical groups show halting or even partial decrease in *Aβ* deposition level from baseline to follow-up images after 24 months. Halting or decrease is considered as the overall brain deposition. This group comprises 16 AD, 20 LMCI, 9 MCI, 29 EMCI, and 26 CN. This effect is likely bound to dynamics of AD (*11*), imprecise diagnosis as many of them were MCI, or non-standard treatments which the patients underwent and might have halted or reduced the proteins deposition (*24*). Indeed, many subjects were of old age taking different types of supplements (as ginkgo biloba and multivitamins) in heterogeneous manner and some Levetiracetam, or other drugs for reasons unrelated to neurodegeneration. Investigating possible effects of those supplements and drugs is beyond the purpose of the paper, and very challenging given all confounding factors. We are instead interested in investigating the capacity of the trained model to handle cases with halting and decreasing *Aβ* deposition, in a given heterogeneous context or possible misdiagnosis.

In table S1, we compare the results on the decreasing subjects using the configurations with the best accuracy on the whole dataset (for ESM, the first configurations with all parameters set to 0). Notably, the MAR model ran on all subjects, and then we considered errors on the decreasing ones. Performing training only on the decreasing group produced worse results, with an MSE of 0.0055 and a PCC of 0.9182, while the GCN model showed better performance training only on decreasing subjects than on the whole dataset, for which it would manifest an MSE of 0.0639 and a PCC of 0.8763. This phenomenon could suggest that patterns of decreasing and increasing subjects are not disjoint, but only the MAR model can capture them training on a larger cohort, gaining better insights on such mechanisms. In figure S2, we analyzed the regional errors of these models on decreasing subjects.

## Discussion

In this study we investigated the capacity of predicting future misfolded *Aβ* from a baseline amyloid PET scan. We will discuss here performance findings in the context of the clinical relevance for the neurodegenerative field.

Surprisingly, the GCN model, despite the natural application of graph neural networks to this context, did not provide remarkable results. A possible explanation can be an insufficient amount of data.

As shown in figure S1, regional errors of all models show consistent values in correspondence to regions as *nucleus reuniens, ventral tegmental area, locus coeruleus, and raphe nucleus (both dorsal and median)*. The Mann-Whitney U test applied to the distributions of regional errors shows the validity of MAR against ESM and GCN, but an equivalence with NDM in this case. Retaining only decreasing subjects shows a different perspective on the models behavior. The MAR model is able to capture the patterns of decrements in regional concentration, especially when trained on the whole dataset, proving a good capacity of generalization. Conversely, ESM and NDM simulative models are built to track exclusively increments, without considering otherwise. This is fundamental in the context of the recent findings regarding *Aβ*, which shifted the focus from a deterministic to a probabilistic approach of AD. This new framework categorizes the disease into three variants: autosomal dominant AD, APOE 4 (the most important genetic risk factor for AD), and APOE 4-unrelated sporadic AD. These variants show different weight of the amyloid cascade and increasing weight of stochastic factors (e.g., environmental, risk genes) (*13*), rather than considering *Aβ* as a sufficient and necessary causative agent. Additionally, AD is characterized by different clinical phenotypes, such as the “language” variant (primary progressive aphasia), the “visual” variant (posterior cortical atrophy), and the “behavioral” variant (*25, 26*). To date, it is still unclear whether different spatio-temporal *Aβ* trajectories are linked to these different variants, in contrast to *τ* PET (*27*). Similarly, it is still unclear how *Aβ* distributions might be related to the AD probabilistic model. *Aβ* deposition begins several years prior to the clinical onset and accumulation (*28*). In this scenario it is not possible to capture directly early differentiation in *Aβ* brain deposition. Here, we provided a new model capable of predicting future *Aβ* deposition. This model could fill the gap with the previous knowledge about the relationship between *Aβ* distribution and AD. Further developments could consider the prediction of. The joint prediction of *Aβ* and *τ* spreading could pave the way to the development of new biomarkers for differential diagnosis. Recently, it has been suggested that limbic-predominant age-related TDP-43 encephalopathy (LATE) might account for 15–20% of all clinically diagnosed AD cases, and although LATE-NC can exist in tandem with other pathologies, most commonly AD-NC (*29*), it is a separate pathology. TDP-43 aggregation in the brain is necessary for LATE-NC diagnosis, with progression moving from the amygdala to the hippocampus to the middle frontal gyrus. By contrast, AD-NC follow a different trajectory. To date, TDP-43 is measurable only on autopsy. Until an in-vivo biomarker is developed, a predictive model using molecular PET scans could help to identify those subjects exhibiting different MP spreading.

Performance of all models peculiarly drop over the MCI clinical group. For MAR and GCN models it would be conceivable that this is due to its size (30 subjects), but also other models follow, pointing to potentially different spreading dynamics.

The results of the MAR for CDR prediction, listed in table 2, point to substantially better performance on CN prediction, which tends to get worse moving towards the AD category. This tendency is in the distribution of predicted CDR scores, shown in figure 3. Seeing the distribution of the true CDRs, it appears that MAR doesn’t have enough data to learn this multimodal distribution.

**Table 1:**
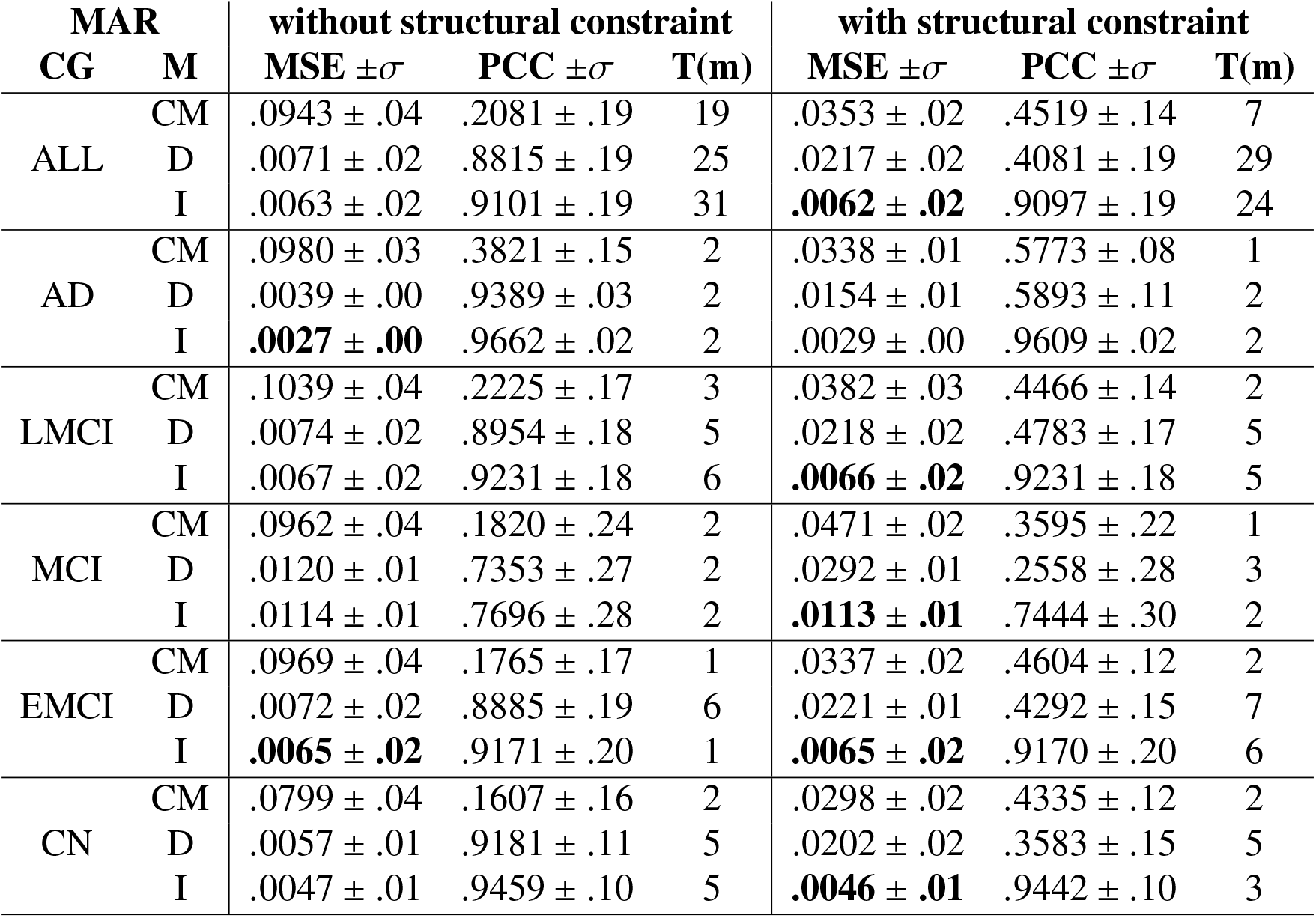
Performance of MAR across all clinical groups (CG) and with several configurations. We report the lower MSE for each CG in bold, preferring a high PCC in case of parity (as in EMCI results). Moreover, this table reports the different results depending on the initialization of the MAR: structural connectivity matrix (CM), diagonal matrix (D), andidentity matrix (I): with and witohut structural constrain.

**Table 2:** Performance of MAR for CDR prediction across all clinical groups (CG). We highlighted in bold the lowest absolute error (AE).

| CDR MAR |  |  |  |  |  |  |  |  |
| --- | --- | --- | --- | --- | --- | --- | --- | --- |
| CG | AE $\pm \sigma$ | T(s) | CG | AE $\pm \sigma$ | T(s) | CG | AE $\pm \sigma$ | T(s) |
| ALL | .3646 ± .36 | 5 | AD | .6500 ± .51 | 1 | LMCI | .3128 ± .26 | 1 |
| MCI | .3200 ± .22 | 1 | EMCI | .2500 ± .24 | 1 | CN | **.0510 ± .15** | 1 |

**Table 3:** Performance of GCN across all clinical groups (CG). We highlighted in bold the lowest MSE in each CG. This table reports the different results depending on the initialization of the GCN: structural connectivity matrix (CM), diagonal matrix (D), andidentity matrix (I).

| GCN |  |  |  |  |  |  |  |  |  |
| --- | --- | --- | --- | --- | --- | --- | --- | --- | --- |
| CG | M | MSE $\pm \sigma$ | PCC $\pm \sigma$ | T(m) | CG | M | MSE $\pm \sigma$ | PCC $\pm \sigma$ | T(m) |
| ALL | CM | **.0659 ± .05** | .8469 ± .19 | 2655 | AD | CM | **.0562 ± .02** | .8496 ± .07 | 38 |
| | D | .0671 ± .05 | .8472 ± .19 | 2628 | | D | .0608 ± .02 | .8617 ± .07 | 37 |
| | I | .0696 ± .05 | .8478 ± .20 | 2672 | | I | .0622 ± .02 | .8554 ± .07 | 37 |
| LMCI | CM | .0744 ± .06 | .8360 ± .18 | 138 | MCI | CM | .0875 ± .04 | .6433 ± .25 | 60 |
| | D | .0752 ± .05 | .8372 ± .18 | 138 | | D | .0881 ± .04 | .6610 ± .25 | 61 |
| | I | **.0724 ± .06** | .8385 ± .18 | 138 | | I | **.0857 ± .04** | .6860 ± .21 | 60 |
| EMCI | CM | .0646 ± .06 | .8423 ± .19 | 228 | CN | CM | .0599 ± .04 | .8600 ± .12 | 172 |
| | D | .0661 ± .06 | .8430 ± .19 | 229 | | D | .0584 ± .04 | .8598 ± .13 | 171 |
| | I | **.0645 ± .06** | .8452 ± .20 | 227 | | I | **.0558 ± .03** | .8665 ± .11 | 168 |

**Table 4:** Performance of ESM across all clinical groups (CG), split by inclusion of Gaussian noise (right) or not (left). Notably, due to our adaptation, the ESM seems insensible to the range of values applied to its parameters [0..1], also because of the small number of optimal iterations.

| ESM | | | $\mu = \sigma = 0$ | | | $\mu = \sigma = 1$ | | |
| --- | --- | --- | --- | --- | --- | --- | --- | --- |
| CG | $\beta_0$ | $\delta_0$ | MSE $\pm \sigma$ | PCC $\pm \sigma$ | T(m) | MSE $\pm \sigma$ | PCC $\pm \sigma$ | T(m) |
| ALL | 0 | 0 | .0065 ± .02 | .9083 ± .19 | 360 | .0065 ± .02 | .9083 ± .19 | 378 |
| | 1 | 1 | .0065 ± .02 | .9083 ± .19 | 383 | .0065 ± .02 | .9083 ± .19 | 378 |
| AD | 0 | 0 | .0040 ± .00 | .9658 ± .02 | 41 | .0040 ± .00 | .9658 ± .02 | 43 |
| | 1 | 1 | .0040 ± .00 | .9658 ± .02 | 43 | .0040 ± .00 | .9658 ± .02 | 43 |
| LMCI | 0 | 0 | .0069 ± .02 | .9216 ± .18 | 80 | .0069 ± .02 | .9215 ± .18 | 84 |
| | 1 | 1 | .0069 ± .02 | .9216 ± .18 | 85 | .0069 ± .02 | .9215 ± .18 | 84 |
| MCI | 0 | 0 | .0105 ± .01 | .7667 ± .29 | 51 | .0105 ± .01 | .7667 ± .29 | 54 |
| | 1 | 1 | .0105 ± .01 | .7667 ± .29 | 54 | .0105 ± .01 | .7667 ± .29 | 54 |
| EMCI | 0 | 0 | .0068 ± .02 | .9152 ± .20 | 102 | .0068 ± .02 | .9083 ± .19 | 107 |
| | 1 | 1 | .0068 ± .02 | .9152 ± .20 | 108 | .0068 ± .02 | .9083 ± .19 | 107 |
| CN | 0 | 0 | .0046 ± .01 | .9443 ± .10 | 87 | .0046 ± .01 | .9442 ± .10 | 91 |
| | 1 | 1 | .0046 ± .01 | .9443 ± .10 | 92 | .0046 ± .01 | .9442 ± .10 | 91 |

**Table 5:** Performance of NDM across all clinical groups (CG), for each value of *β*. The lower MSE for each CG is in bold.

| NDM<br>CG | $\beta$ | MSE $\pm \sigma$ | PCC $\pm \sigma$ | T(m) | $\beta$ | MSE $\pm \sigma$ | PCC $\pm \sigma$ | T(m) |
| --- | --- | --- | --- | --- | --- | --- | --- | --- |
| ALL | .0 | .0063 ± .01 | .90 ± .16 | 173 | .1 | .0063 ± .01 | .9044 ± .16 | 163 |
| | .2 | **.0063** ± **.02** | .9100 ± .19 | 162 | .4 | .0063 ± .02 | .9099 ± .19 | 167 |
| | .8 | .0063 ± .02 | .9098 ± .19 | 176 | 1 | .0063 ± .02 | .9098 ± .19 | 170 |
| AD | .0 | .0036 ± .00 | .9663 ± .02 | 20 | .1 | .0036 ± .00 | .9662 ± .02 | 18 |
| | .2 | .0036 ± .00 | .9661 ± .02 | 18 | .4 | .0036 ± .00 | .9660 ± .02 | 19 |
| | .8 | **.0035** ± **.00** | .9657 ± .02 | 20 | 1 | .0035 ± .00 | .9656 ± .02 | 19 |
| LMCI | .0 | **.0066** ± **.02** | .9230 ± .02 | 38 | .1 | .0066 ± .02 | .9229 ± .18 | 36 |
| | .2 | .0066 ± .02 | .9229 ± .18 | 36 | .4 | .0066 ± .00 | .9228 ± .18 | 37 |
| | .8 | .0066 ± .02 | .9226 ± .18 | 39 | 1 | .0066 ± .02 | .9225 ± .18 | 38 |
| MCI | .0 | .0104 ± .01 | .7701 ± .28 | 25 | .1 | .0104 ± .01 | .7702 ± .28 | 23 |
| | .2 | .0104 ± .01 | .7703 ± .28 | 23 | .4 | .0104 ± .01 | .7704 ± .28 | 24 |
| | .8 | .0104 ± .01 | .7706 ± .28 | 25 | 1 | **.0104** ± **.01** | .7707 ± .28 | 24 |
| EMCI | .0 | .0066 ± .02 | .9169 ± .20 | 49 | .1 | .0066 ± .02 | .9169 ± .20 | 46 |
| | .2 | .0066 ± .02 | .9169 ± .20 | 46 | .4 | .0066 ± .02 | .9168 ± .20 | 47 |
| | .8 | .0066 ± .02 | .9167 ± .20 | 50 | 1 | **.0065** ± **.02** | .9167 ± .20 | 48 |
| CN | .0 | .0045 ± .01 | .9457 ± .10 | 42 | .1 | .0045 ± .01 | .9457 ± .10 | 39 |
| | .2 | .0045 ± .01 | .9457 ± .10 | 39 | .4 | .0045 ± .01 | .9456 ± .10 | 40 |
| | .8 | **.0044** ± **.01** | .9454 ± .10 | 42 | 1 | **.0044** ± **.01** | .9454 ± .10 | 41 |

The mechanisms by which misfolded proteins spread through the brain remains controversial. In the current work we did not attempt to answer whether the Braak hypothesis (*30*) was valid within our experiments. The presence of both decreasing and increasing *Aβ* confirms the “two-stage” model. In the two-stage model, increased early *Aβ* accumulation occurs. In the second this change subsequently hastens the spreading of *τ*. Our model can predict the spreading independently on the clinical stage and could be used to plan specific therapies depending on the spatial distribution of MP deposition. This is extremely important to design both preventive actions and rehabilitative protocols, such as non-invasive brain stimulation interventions (NIBS). In the last years, NIBS emerged as a promising approach to restore cognitive functions impaired in neurodegenerative disorders. However, previous studies relied on a “one-size fits-all” approach (for a review see (*31*)). The clinical effectiveness induced by NIBS may be maximized by designing personalized interventions (*32*). Our model could help to guide the implementation of new personalized NIBS approaches, tailoring the parameters and location of stimulation (e.g., target regions) to the individualized pattern of MP propagation computed from a baseline PET scan. The advantage of this approach relies on the identification of those brain regions which will display the largest amount of MP accumulation. Thus, future studies applying our model would assess whether stimulation of these regions would maximize NIBS clinical efficacy and the maintenance of the results, in line with the assumption that NIBS protocol applied in the earliest stages could be more effective (*33*).

Most of previous studies (*14, 15*) capitalize on simulating protein spreading over a long period of time on an healthy connectome. However, given neurodegeneration it is inevitable that connectome of AD patients and healthy control will be different. In this study we assume that a more reliable approach is given by using the individual connectome. As a limitation, we do not consider further changes in the connectome which will inevitably happen with the progression of disease (*34*), which is notwithstanding also neglected by the other approaches simulation-based. In our study we have not been able to understand what are the specific patterns that discriminate with one time point whether the trend of *Aβ* is decreasing or increasing and therefore identifying the phase, though the algorithm is able to perform this prediction.

Lastly, studies highlighted different trajectories and characterization of AD. Temporal trajectory models illustrate that there is a distinct progression pattern of AD especially if taking into account sex and APOE genotyping (*35*). Furthermore, limbic-predominant and medial temporal lobe-sparing patterns, also highlight possible temporal patterns of atypical clinical variants of AD (*36*). These aspects appear to be yet open discussions, with studies hypothetizing that subtypes might be just different stages of the disease, urging for clearer longitudinal analysis (*37,38*), while other relatively confirming the sub-typing via longitudinal analysis (*39*). Given the complexity and the available data, investigation of different trajectories by using the proposed model is a future work.

Nevertheless, a better understanding of brain connectivity, jointly to misfolded protein deposition prediction, could open the door to brand-new, specifically designed network therapies, guided by knowledge of networks important in various clinical phenotypes. Moreover, new insights about the relationship between macro-scale systems, as the one presented in this work, and micro-scale molecular processes might help to identify which individuals would benefit most.

Through our experiments we assessed the validity and the characteristics of MAR in a novel context, showing the potential to become a fast and reliable model for instantaneous predictions. We trusted this potential realizing an online platform able to perform such predictions starting from a pre-processed or raw PET image and displaying all necessary information to clinicians or other researchers.

## Methods

### Selection of subjects

From the ADNI database, we selected DWI, T1, and PET suitable images, converting them from DICOM to NIFTI using the program *dcm2niix* (*40*) following BIDS format (https://bids.neuroimaging.io/).

We considered T1 images acquired with 3-Tesla scanners and belonging to sagittal IR-SPGR (Inversion Recovery Spoiled Gradient Recalled) or MPRAGE (magnetization-prepared 180 degrees radio-frequency pulses and rapid gradient-echo) protocols.

We selected DWI images having at least 30 volumes and an axial acquisition on 3-Tesla scanners, with a posterior-to-anterior acquisition. We extracted from multi-shell (127 volumes, b={0,500,1000,2000}s/mm^2^) images the volumes and b-vectors corresponding to single-shell values (b={0,1000}s/mm^2^), obtaining 50 single-shell volumes for each. We kept only the first b-values equal to 0 s/mm^2^, not those following greater values. The choice of extracting singleshell values has been driven by the greater compatibility with preprocessing, harmonization, and tractography algorithms, as shown later.

We selected PET images obtained with AV45, FBB, and PIB tracers, at 2 time points distanced 24 months, correcting the space when not in the standard one with the command *fslreorient2std*, and we occasionally discarded the first frames when they were of poor quality through visual assessment.

We merged suitable images as depicted in figure S3, resulting in 212 individuals (24 AD, 47 LMCI, 30 MCI, 60 EMCI, and 51 CN).

### Preprocessing

Each image went through a dedicated pipeline to facilitate the further steps, written in Python programming language and relying on FSL (*41*), Dipy (*42*), creating a workflow using Nipype (*43*) encapsulation.

For T1 images, we performed brain extraction using BET2, then we registered to the Automated Anatomical Labeling 3 (AAL3) atlas (2mm resolution) (*44*) through FLIRT and performed brain segmentation utilizing FAST.

For DWI, we performed a slightly different brain extraction with BET2, proceeding with LPCA denoising and Gibbs correction from Dipy, then Eddy correction through the homonym component in FSL, and then co-registered to the previously registered T1 image. For Eddy correction and registration, both implying a roto-translation, b-vectors have been modified accordingly to maintain a correct gradient table, as pointed out by (*45*), using Dipy.

For PET, we performed motion correction through a co-registration of frames using Dipy, then we performed an average of these frames, obviously across time, using fslmaths, then registered to a reference volume of AAL3 with skull (1mm resolution). The standard resolution AAL3 atlas used for T1 registration helps with memory and time consumption during tractography, which doesn’t require high precision to infer long-range connections between brain regions. Conversely, PET registration required a reference volume with a skull, available only for the 1mm resolution, which allowed greater accuracy in allocating Amyloid-*β* concentrations, and then better predictions.

### Harmonization

Different scanners can induce distinct artifacts in the DWI images acquired, instilling noise and potential hindrances in their manipulation. (*46–48*) depicted and addressed this problem, expressing the need for a harmonization of images from different scanners. We gathered together images depending on their volumes (31, 36, 46, 50, 54, or 55), and we harmonized them against the 46 (the most numerous) through a python pipeline (*49*), which required the extraction of single-shells volumes from multi-shell images as depicted above.

### Transformation of PET concentrations

After preprocessing, we accessed the PET images of each subject, normalized them by the maximum value of the image in the range [0, 1], and computed the average regional concentrations for each image following the 166 ROIs of AAL3. We averaged again for each category and applied a z-score normalization against the baseline PET concentrations for CN subjects as done in (*15*). Then, z-scores were remapped into a [0, 1] interval using a standard logistic function, obtaining values displayed in figure 5.

**Figure 5:**
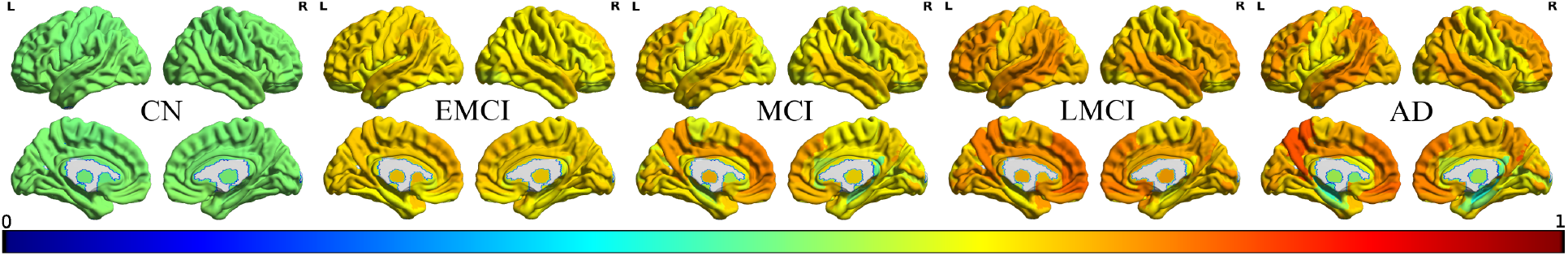
A graphical representation of Amyloid-*β* concentrations for each clinical stage, normalized for the CN regional values (left) and ignoring voxels not contained in the AAL3 atlas (gray).

### Tractography

After the harmonization stage, we used DWI and T1 images to infer white matter fibers using methods provided by Dipy. Firstly, we binarized seeds value, then obtained the response function distribution and applied a constrained spherical deconvolution model of the 6th spherical harmonic order. The fitted model has been used to generate probabilistic streamline distribution following the anatomically-constrained tractography (ACT) stopping criterion, which uses tissues segmented during the preprocessing of T1 images to handle and stop tracking of fibers (*50*). The angle threshold was 20 degrees, and the imposed minimum length of streamlines was 10, generally reputed noise (*51*).

We matched the remaining streamlines to the AAL3 atlas labels and kept inter-regional connections, creating a connectivity matrix (CM), erasing its diagonal, and normalizing it for its maximum value. We specify that this is the standard practice for CM normalization, but in simulations scenarios, we put ones on the main diagonal of CM. Notably, the AAL3 atlas has up to 170 labels, of which four are empty to maintain compatibility (35, 36, 81, and 82) with previous atlas versions. We dropped them to obtain the 166 effective regions used for predictions.

### CDR

From ADNI repository we retrieved the total score of the CDR scale. This clinical questionnaire investigated daily function in the domains of memory, orientation, judgment and problem solving, community affairs, home and hobbies, and personal care from information collected from the informant interview as well as the individual interview. When the global scores were not available, we computed them from this six subcategories following the ADNI criteria available in ADNI procedures.

### Common simulation settings

We computed, for each simulation, the average MSE and PCC across folds, plus their standard deviation. We also saved the time necessary for the simulation and the number of threads used to parallelize the computation. A list of subjects, expanded with MSE and PCC in both the training and testing phase, is returned to provide insights into the quality of individual predictions. Furthermore, we compute a list of average regional errors to understand strong and weak ROIs for each model.

### MAR

The MAR model defines a linear dependency between the previous values of a set of variables and their next state, as stated in the equation (1), showing potential issues in case of a large number of regions or noise (*21*). This drawback drove us to adopt the reasonable number of ROIs of the AAL3 atlas. The term *multivariate* indicates a modeling of the outcome based on longitudinal (multiple time points for the same subject) or clustered data (*52*). Given the limited amount of time points and subjects available with PET data, we opted for the second approach, enabling a more fair comparison with other predictive models which don’t account for multiple time points and a more accurate analysis of the stage of disease with better scores for each model.

Given the values of *A* − *β* at two different time points 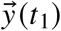 and 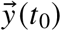, the approach stems from the definition of reconstruction error 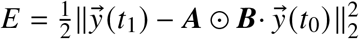. The dimension of the matrix *A* is 166*x*166 and both 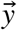 are row vectors of length 166. The reconstruction is minimized by the optimal matrix *Â*, obtained deriving the error according to *A* as in Equation (1).

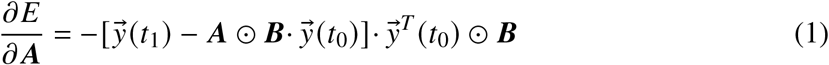

This formulation is a first-order model, based on two time points, described in (*21*), where the Hadamard (element-wise) product ⊙ with the indicator matrix ***B***, representing the structural bias, constrains the model to use the anatomical connectivity, and the starting point of the optimization process is variable. The considered values for the A matrix were the connectivity matrix, a diagonal matrix containing the *A* − *β* concentration at baseline, or a simple identity matrix. In all cases, A has dimensions 166×166. If we used the CM, we set its diagonal values to 1 to consider endogenous raising in concentrations.

***B*** is a binary matrix containing ones in the position of existing structural connections and zeroes for others. We investigate its role in providing faster or more anatomically feasible solutions, measuring performance with and without it.

The optimization is a classical gradient-descent approach with the static update rule *A* = (*A* + *η*· ∂*E*/∂*A*), applied using an *η* corresponding to 1*e*^−5^. The termination criterion was met when 3 million iterations were completed or when an increment of the value of the reconstruction error *E* was registered. We tested this approach through a leave-one-out cross-validation for each category and whole dataset. We averaged the coefficient matrices of all subjects within the training set as in *average ensembling* technique before testing against the corresponding test set for that fold.

MAR predicted CDR scores using the slightly different gradient function in Equation (2). In this equation, 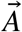 is a vector 166 regions long, each element initialized as one plus the CDR at baseline. The gradient descent halted at the first increment of the error function.

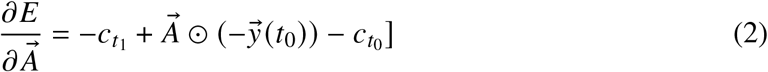

The prediction is computed as 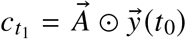, using all the trained 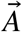 vectors in an ensemble, which notably is the mean CDR score predicted, due to the linearity of the model. The absolute error is then kept and used to measure the goodness of the prediction.

### GCN

Due to the peculiar structure of our data, we adopted a neural network architecture highly specialized in graph learning and able to transpose the concept of *convolution*, traditionally bound to images, to these structures presenting variable neighborhoods. In Equation (3) we present the forward propagation rule for convolution.

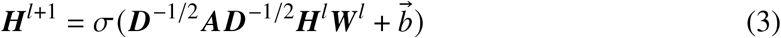

Where ***H***^*l*^ is the matrix of activation for the *l*_*th*_ layer, employing the rectified linear unit (ReLU) activation function *σ* on the degree matrix ***D***, and on the weight matrix ***W***^*l*^ for the *l*_*th*_ layer. The adjacency matrix A allows learning of features from surrounding nodes.

The convolution consisted of two layers interleaved with batch normalization, flatten, and dropout layers. The batch normalization layer provided by the Keras library remapped the input of shape (8, 166, 32) to have a mean close to 0 and a standard deviation of 1. The flatten layer reshaped the input in (8, 5312), and the dropout layer iteratively deactivated 30% of the available nodes along the training phase, making the model more robust, simulating different architectural configurations, and less prone to overfitting showing the smaller amount of samples to each node.

Two dense layers followed the convolution, the first with 32 nodes and a tanh activation function and the second with 166 nodes and a linear activation function.

The training executed for 20’000 epochs using Adam algorithm with *η* equal to 1*e*^−5^ for gradient descent. We assessed the performance using the leave-one-out cross-validation approach.

### ESM

The ESM conceptualizes the spreading of misfolded proteins as an epidemic process, therefore it is based on stochastic infection and clearance of MP, represented as 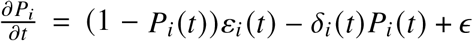.

The authors adopted the Gini coefficient to prevent excessive dispersions of misfolded proteins, which would cause steep decrements in local concentrations. *P*_*i*_ describes the probability for the *i*_*th*_ region being infested at time *t*, dependently on *ε*_*i*_ (*t*), which holds the chance of receiving MP agents, *δ*_*i*_ (*t*), which is the probability of being clean of MP at time *t*, and *ϵ*, which represents Gaussian noise related to the stochastic processes, with mean *µ* and standard deviation *σ*. We initialized the vector of probabilities *P* with ones corresponding to regions where there were *A* − *β* and zeroes elsewhere. As reported by the same authors (*14*), the simulation sits on four unknown constant parameters [*β*_0_, *δ*_0_, *µ, σ*].

Despite how originally formulated in (*14*), we chose to predict the diffusion *in the future* instead of reconstructing the past of an AD subject. To accomplish this objective, we added a further step formulated in equation (4), where ***C*** is the connectivity matrix.

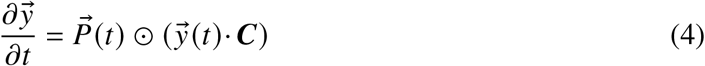

The simulations parameters comprised a *η* of 1*e*^−2^ and two iterations empirically estimated as the average optimal amount for each subject. We set a larger *η* for the many hyper-parameters smaller than one, which could lead to possible underflows and inconsistent variations in the concentrations. The assessment of iterations has been necessary to assure a fair comparison with learning models, and due to the lack of authors’ specifications about this and other parameters obtained in (*14*). Therefore, even [*β*_0_, *δ*_0_, *µ, σ*] impact has been tested.

### NDM

The NDM exploits a diffusion kernel to model a rate of diffusion, considered constant in time, through the normalized Laplacian of the connectivity matrix, which decomposes in the form 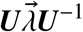, with ***U*** and 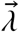 corresponding to eigenvectors and eigenvalues, respectively. In (*15*), the relationship between gray matter atrophy and hypo-metabolism of glucose was considered to predict future atrophy. We applied the same equation to *A* − *β* diffusion as 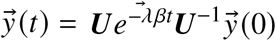, where *β* is a constant rate of network diffusion, and *t* is the prediction time.

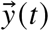 is the vector containing regional concentrations of *A* − *β* at time *t. λ*_*i*_ is the *i*_*th*_ eigenvalue of the Laplacian of the connectivity matrix. Notably, eigenvectors have been added with 1*e*^−10^ to avoid divisions by zero.

We transposed this method to *A* − *β* prediction and assessed the optimal average iterations for ESM, found to be 2318 for our dataset, and an evaluation of the optimal value of *β* across each category and the whole cohort. We applied a time step of 1*e*^−5^.

## Acknowledgments

This publication is supported by the European Union’s Horizon 2020 research and innovation programme under grant agreement Sano No 857533. This publication is supported by Sano project carried out within the International Research Agendas programme of the Foundation for Polish Science, co-financed by the European Union under the European Regional Development Fund. This research was supported in part by the PLGrid infrastructure on the Prometheus computer cluster (kdm.cyfronet.pl/portal/Prometheus:en). The authors gratefully acknowledge the computing time provided on the JURECA supercomputer at Forschungszentrum Jülich (*53*). We wish to express sincere gratitude to our colleague Joan Falco Roget for his valuable insights and feedback on statistical and mathematical methods here applied.

Data collection and sharing for this project was funded by the Alzheimer’s Disease Neuroimaging Initiative (ADNI) (National Institutes of Health Grant U01 AG024904) and DOD ADNI (Department of Defense award number W81XWH-12-2-0012). ADNI is funded by the National Institute on Aging, the National Institute of Biomedical Imaging and Bioengineering, and through generous contributions from the following: AbbVie, Alzheimer’s Association; Alzheimer’s Drug Discovery Foundation; Araclon Biotech; BioClinica, Inc.; Biogen; Bristol-Myers Squibb Company; CereSpir, Inc.; Cogstate; Eisai Inc.; Elan Pharmaceuticals, Inc.; Eli Lilly and Company; EuroImmun; F. Hoffmann-La Roche Ltd and its affiliated company Genentech, Inc.; Fujirebio; GE Healthcare; IXICO Ltd.;Janssen Alzheimer Immunotherapy Research Development, LLC.; Johnson & Johnson Pharmaceutical Research Development LLC.; Lumosity; Lundbeck; Merck Co., Inc.;Meso Scale Diagnostics, LLC.; NeuroRx Research; Neurotrack Technologies; Novartis Pharmaceuticals Corporation; Pfizer Inc.; Piramal Imaging; Servier; Takeda Pharmaceutical Company; and Transition Therapeutics. The Canadian Institutes of Health Research is providing funds to support ADNI clinical sites in Canada. Private sector contributions are facilitated by the Foundation for the National Institutes of Health (www.fnih.org). The grantee organization is the Northern California Institute for Research and Education, and the study is coordinated by the Alzheimer’s Therapeutic Research Institute at the University of Southern California. ADNI data are disseminated by the Laboratory for Neuro Imaging at the University of Southern California.

## Data and Code availability

All code used to produce the results presented in this paper are available at the URL https://github.com/LucaGherardini/PredictionMisfoldedProteins.

## Supplementary material

**Figure S1:**
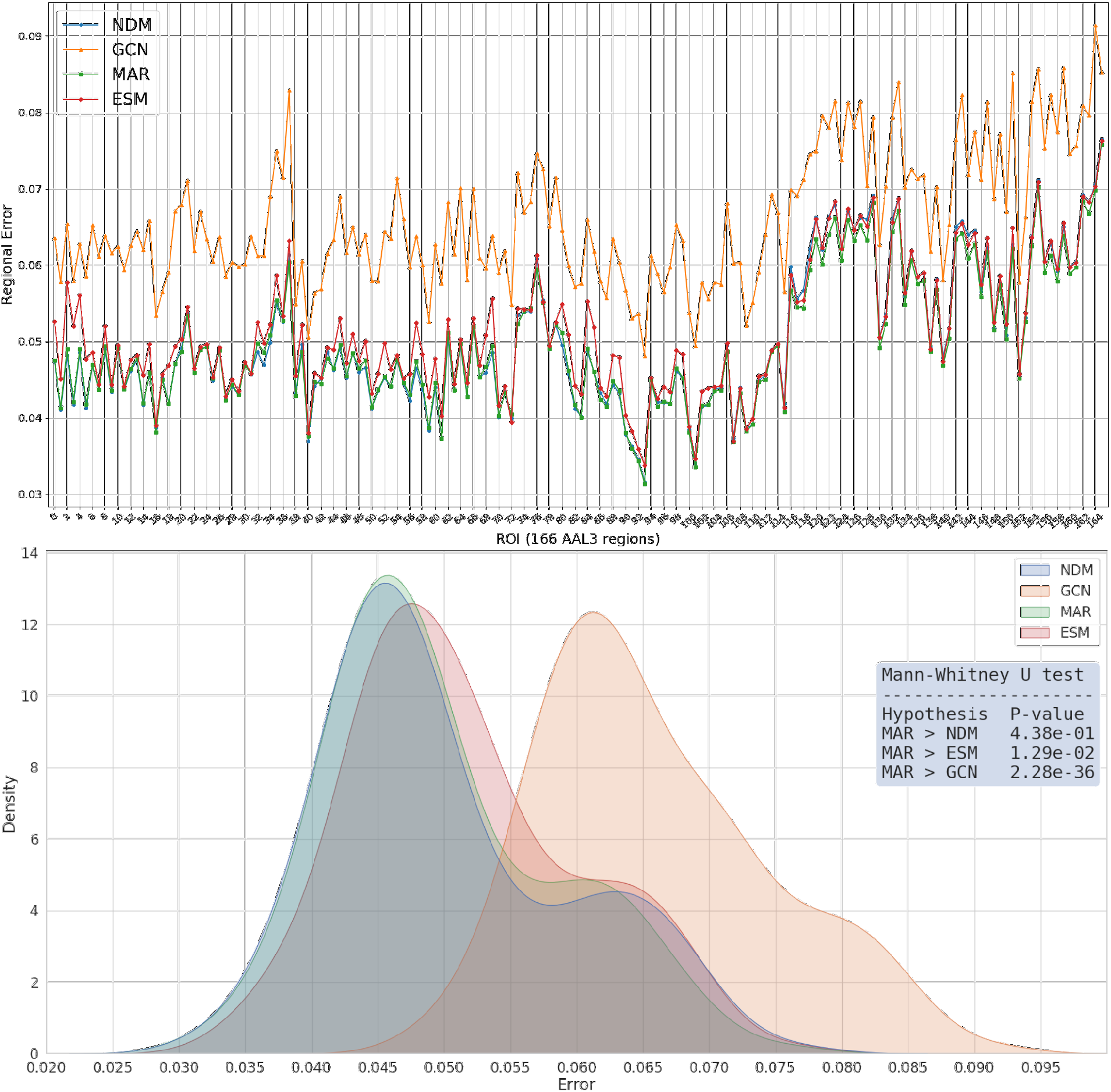
We display, for each model, regional errors (top) and relative error density (bottom). In the plot above, errors of MAR and NDM follow an overlapping trajectory under those of ESM and GCN. Notably, the GCN model has substantially higher errors, and therefore its error density is shifted to the right. The ESM and NDM densities are slightly more right-skewed than MAR, revealing sporadic higher errors. The Mann-Whitney U test (in box) reveals smaller errors in MAR than in ESM and GCN, but this result is not strongly confirmed for NDM, against which errors are not strictly smaller.

**Figure S2:**
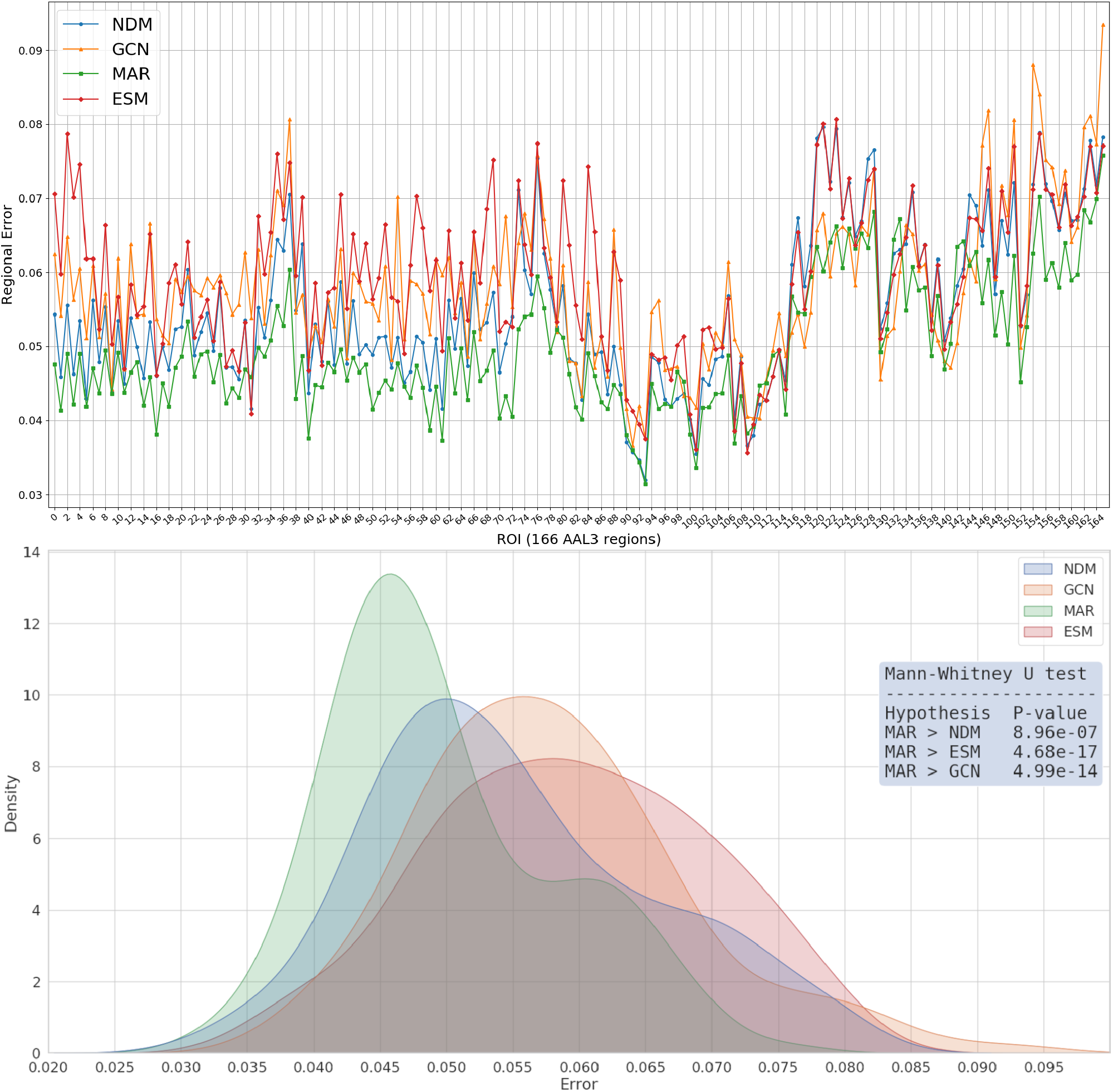
We display, specularly to figure S1, regional errors for decreasing subjects. Here a neat gap between MAR and other models appears, and is confirmed by the Mann-Whitney U test.

**Figure S3:**
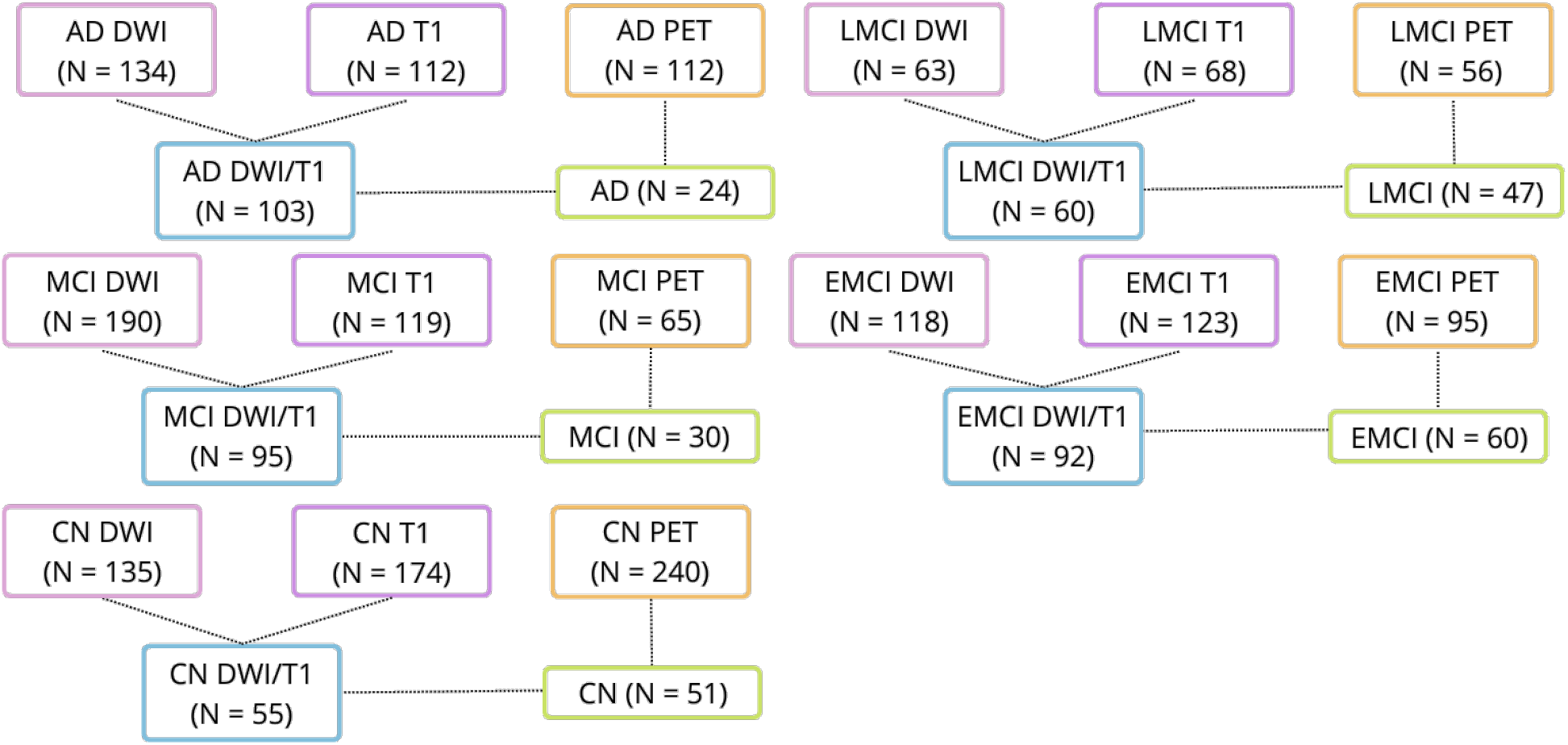
The selection process of suitable subjects, with the amount for each image specified between parentheses.

**Figure S4:**
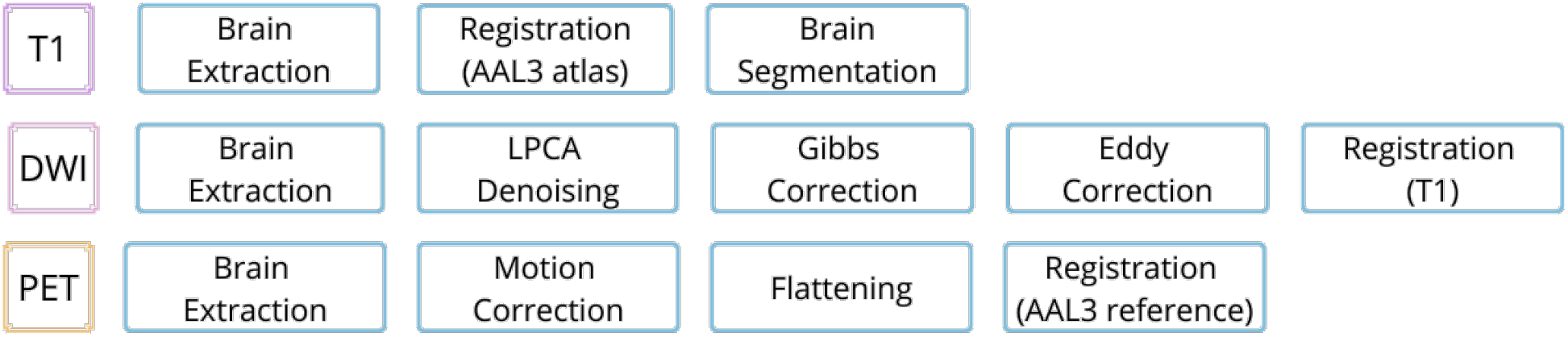
Preprocessing steps for each kind of image. Notably, *Eddy Correction* and *Registration to T1* in DWI pathway required correction of b-vectors.

**Table S1:** Performance of all models on decreasing subjects, using the best configurations for the whole dataset (for the ESM, the first one). For MAR, training exclusively on the decreasing subjects produced worse results, therefore we report the density of regional errors on the whole dataset.

| Decreasing subjects |  |  |  |  |  |  |  |
| --- | --- | --- | --- | --- | --- | --- | --- |
| Model | MSE $\pm \sigma$ | PCC $\pm \sigma$ | T(m) | Model | MSE $\pm \sigma$ | PCC $\pm \sigma$ | T(m) |
| MAR | .0039 ± .01 | .9479 ± .09 | 102 | GCN | .0579 ± .05 | .8776 ± .18 | 63 |
| ESM | .0081 ± .02 | .9336 ± .16 | 155 | NDM | .0075 ± .02 | .9342 ± .16 | 71 |

